# Vasopressin V2 is a promiscuous G protein-coupled receptor that is biased by its peptide ligands

**DOI:** 10.1101/2021.01.28.427950

**Authors:** Franziska M. Heydenreich, Bianca Plouffe, Aurélien Rizk, Dalibor Milić, Joris Zhou, Billy Breton, Christian Le Gouill, Asuka Inoue, Michel Bouvier, Dmitry B. Veprintsev

## Abstract

Activation of the G protein-coupled receptors by agonists may result in the activation of one or more G proteins, and in the recruitment of arrestins. The balance of activation of different pathways can be influenced by the ligand. Using BRET-based biosensors, we showed that the vasopressin V2 receptor activates or at least engages many different G proteins across all G protein subfamilies in response to its native agonist arginine vasopressin (AVP). This includes members of the Gi/o and G12/13 families that have not been previously reported. These signalling pathways are also activated by the synthetic peptide desmopressin and natural homologs of AVP, namely oxytocin and the non-mammalian hormone vasotocin. They demonstrated varying degrees of functional selectivity relative to AVP, as quantified using the operational model for quantifying ligand bias. Additionally, we modelled G protein activation as a Michaelis-Menten reaction. This approach provided a complementary way to quantify signalling bias, with an added benefit of clear separation of the effects of ligand affinity from the intrinsic activity of the receptor. These results showed that V2 receptor is not only promiscuous in its ability to engage several G proteins, but also that its signalling profile could be easily biased by small structural changes in the ligand.

## Introduction

G protein-coupled receptors (GPCRs) are a family of membrane proteins involved in many physiological processes including vision, smell, hormone regulation and neurotransmission. Their extracellular ligand-binding sites and their impact on cellular signalling make them prime drug targets [1]. GPCRs translate ligand-binding events into cellular signals via activation of heterotrimeric G proteins and arrestins. Despite a large variety of incoming signals, there are only 16 different Gα, 5 Gβ and 12 Gγ subunits [2, 3], two β-arrestins and two visual arrestins mediating intracellular signalling. Experimental evidence shows that many receptors are capable of activating or engaging more than one G protein isoform, not only within but also across the Gs/olf, Gi/o, Gq/11 and G12/13 subfamilies of heterotrimeric G proteins [4–11]. The realisation that some ligands can be agonists for one pathway and antagonists for another led to the development of the concept of biased signalling [12–17]. Importantly, such biased ligands are very promising pharmaceuticals because pharmacological benefits are often associated with one particular pathway while the undesired side-effects are mediated by another [13, 18–22]. To compare ligands with each other, a quantitative measure of their signalling efficacy towards individual effectors and their signalling bias is required.

Several approaches to quantify signalling bias have been suggested (reviewed in [21, 23]) the most widely used approach is based on the Black-Leff operational model [24–26]. This formalism describes the ligand binding and effector output of the receptors and provided a framework for the development of quantitative pharmacology [24]. One of the important aspects of this model is its simplicity while keeping the ability to capture important aspects of the signalling process. On the other hand, it is a heuristic model that links the signalling input (ligand binding) to the signalling output (response), without considering the underlying mechanisms. Alternatively, the process of the G protein activation can be presented as an enzymatic reaction, in which the receptor is an enzyme which catalyses the nucleotide exchange in the G protein. This model is very insightful but as it considers G protein recruitment and nucleotide exchange, it is rather complex and contains multiple parameters. As our understanding of the molecular mechanisms involved in the receptor signalling and desensitisation improved, even more detailed descriptions of the signalling processes using quantitative systems pharmacology approaches have been developed [27]. These approaches hold a lot of promise, but the complexity of the signalling and the corresponding models requires a significant amount of diverse data to be collected, limiting its application for routine screening of compounds for bias. A model with a level of simplicity comparable to the Black-Leff formalism that would allow complementary quantification of the physico-chemical ligand efficacy would help to better classified compounds and understand the mechanistic basis for differential G protein activation.

Here, we show that the V2 receptor can activate or engage many G proteins in addition to the previously reported Gs and Gq, including members of the Gi/o subfamily (Gαi1, Gαi2, Gαi3, Gαz), all members of Gq/11 (Gαq, Gα11, Gα14, Gα15), and G12/13 (Gα12 and Gα13). We compared four closely related natural and synthetic peptide ligands for V2 receptor: AVP, arginine-vasotocin (a non-mammalian analogue of vasopressin, referred to as vasotocin thereafter), oxytocin and the clinically used vasopressin analogue desmopressin. These peptides differ in only one or two amino acids. As these peptides have widely different affinity towards the receptor that skews the bias factor calculations using the operational model, we developed a simple Michaelis-Menten formalism of G protein activation that can better separate the effects of ligand affinity vs efficacy in the calculation of the signalling bias. Overall, these results suggest that even relatively minor structural changes in the ligand can induce significant signalling bias at the V2 receptor.

## Results

### Vasopressin V2 receptor recruits members of all G protein families and both β-arrestins

We used biosensors based on bioluminescence resonance energy transfer (BRET) to study engagement of different G proteins. The Gα subunit of the heterotrimeric G protein was tagged with a mutated variant of *Renilla reniformis* luciferase RlucII [28] and the Gγ subunit was N-terminally fused to GFP10 (Fig. 1A). At saturating concentrations of vasopressin, the ligand-mediated ΔBRET obtained with several different Gα proteins fused to RlucII was measured. Vasopressin V2 receptor was able to engage Gαs, Gαi1, Gαi2, Gαi3, Gαz, Gαq, Gα12 and Gα13 to a different extent in presence of the natural ligand, AVP, but failed to recruit or activate GαoA and GαoB (Fig. 1B). This strongly indicates that the V2 receptor can recruit a broader spectrum of G proteins beyond the previously reported Gs and Gq [10, 29]. To investigate V2R coupling specificity towards the members of the Gq/11 family, we used protein kinase C (PKC) biosensor [30] in Gq/11/12/13 knock-out cells [29]. This biosensor reports Gq/11-family mediated activation of protein kinase C biosensor that reports on phosphorylation-induced association of forkhead-associated domains selectively binding to PKC-phosphorylated sites (Fig. 1C). Upon addition of AVP, Gq/11/12/13-deficient HEK293 cells showed PKC activation when complemented with Gαq, Gα11, Gα14 or Gα15 but not in absence of co-expressed Gαq family subunits (Fig. 1D). The specific activation of PKC through co-transfected Gq family members shows that V2R not only couples to but also activates all members of the Gq/11 family. Similarly, previously published cAMP activation data have shown that the V2R activates Gs [10]. Using the Gα-Gγ biosensor (Fig. 1A) we also tested the change in BRET signal for all recruited Gα subunits in combination with three different Gγ subunits: Gγ1, Gγ2 and Gγ5 (Fig. 1E). We found that all combinations except those of Gα12 with Gγ2 and Gγ5 lead to a decrease in BRET signal. This may point towards a difference in the interaction of V2R with Gα12 compared to other Gα subunits and agrees well with a recent study that reported unproductive complex formation between Gα12 and the V2R [31]. In addition to its G protein coupling, the V2R has previously been reported to bind β-arrestin 1 and 2 equally well [32]. We measured β-arrestin recruitment using a recently developed enhanced bystander BRET (ebBRET)-based biosensor that uses RlucII-β-arrestin 1 or 2 and a *Renilla* GFP (rGFP)-tagged CAAX box domain from KRas, which is inserted into the membrane (Fig. 1F) [33]. V2R recruited both β*-*arrestins to the same extent (Fig. 1G) at saturating concentrations of AVP, consistent with its classification as Class B for arrestin recruitment [32].

**Figure 1.**
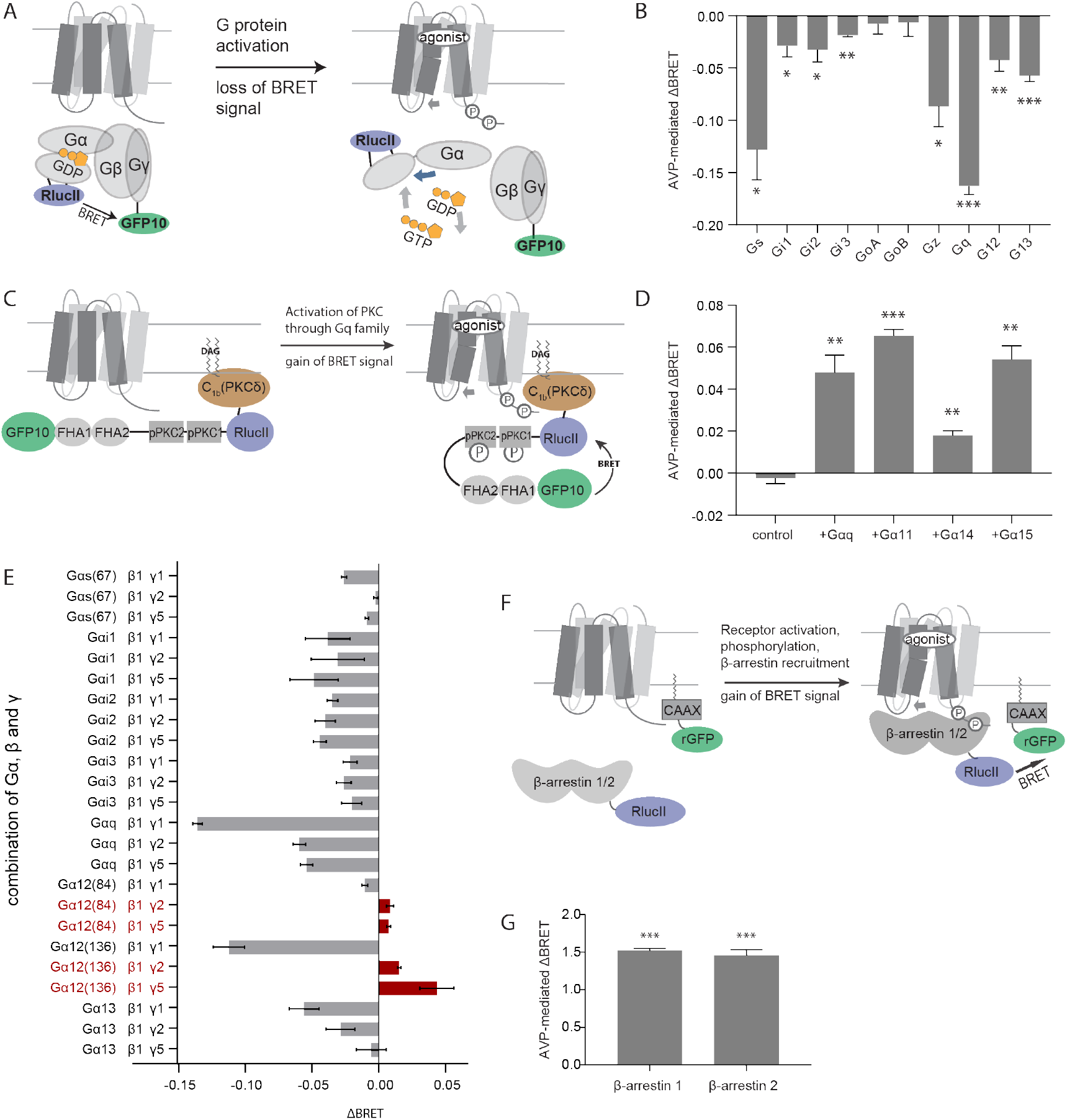
Vasopressin V2 receptor activates Gs/olf, Gi/o, Gq/11 and G_12/13_ family proteins and both β-arrestins. **A.** Schematic overview of the direct G protein BRET-based biosensor. Activation of the heterotrimeric G protein by the GPCR leads to dissociation of the Gα from Gβγ and a conformational change in the Gα domain, which result in a decreased BRET signal. **B.** Overview of the arginine-vasopressin (AVP)-induced change in BRET signal for RlucII-tagged Gα subunits and GFP10-tagged Gγ1. **C.** Schematic overview of the protein kinase C (PKC) biosensor. GFP10 is followed by two phospho-sensing domains, FHA1 and FHA2 and two phospho-PKC (pPKC) sequences which can be phosphorylated by natively-expressed PKC, the RlucII and the C1b domain from PKCδ which binds DAG, leading to membrane recruitment. Activation of the GPCR leads to activation of phospholipase Cβ, followed by accumulation of diacylglycerol (DAG) which activates PKC. The PKC natively expressed in HEK293 cells phosphorylates pPKC1 and 2 domains of the PKC biosensor which causes a conformational change and BRET increase. Through the C1b domain of PKCδ, the sensor is recruited to DAG in the plasma membrane. **D.** Overview of the AVP-induced change in BRET signal for the PKC biosensor when different isoforms from the Gq/11 family are cotransfected in Gq/11/12/13 null HEK293 cells. **E.** G protein engagement at saturating AVP concentrations varies with Gγ subunit. Different combinations of Gα and Gγ lead to differing ΔBRET values, numbers in brackets indicate the amino acid where RlucII was fused to Gα. For Gαs, the alternative fusion position after amino acid 67 was used here. **F.** Schematic overview of the β-arrestin recruitment biosensor. Activation of the GPCR leads to phosphorylation of the C-terminus of the receptor followed by recruitment of β-arrestin. **G.** Overview of the AVP-induced change in BRET signal for RlucII-tagged β-arrestins and rGFP-tagged CAAX domain of Kras. The statistical significance was assessed by a one-sample t-test compared to 0 with n=3 (*p<0.05, **p<0.01 and ***p<0.001). Error bars are shown as standard error of mean (SEM).

### Small differences in peptide ligand sequences give rise to functional selectivity

For determination of signalling bias among peptide ligands, we compared the signalling of the main endogenous agonist AVP to the vasopressin-analogue desmopressin, the non-mammalian vasopressin analogue vasotocin and the natural agonist oxytocin. The nonapeptides all contain a disulphide bridge and differ in either one or two amino acids (Fig. 2A). We tested the effect of all ligands for Gαs, Gαi2, Gαz, Gαq, Gα12 and Gα13 engagement, protein kinase C activation through Gαq, Gα11, Gα14 or Gα15 and the recruitment of β-arrestin 1 and 2. For each effector and ligand combination, we measured concentration-response curves to determine the half-maximal effective concentration (potency, EC50 or pEC50, respectively) (Fig. 2B). In our experiment, the maximal agonist-induced response did not vary across the different ligands tested; all ligands were full agonists for β-arrestin recruitment and G protein engagement. However, the potencies (pEC50) differed by almost 2.5 orders of magnitude between AVP and oxytocin (Table 1). The four ligands also significantly differ in their affinity with AVP being the best binder (single digit nanomolar), followed by vasotocin and desmopressin (low two-digit nanomolar) and oxytocin (low micromolar) as determined by radioligand binding [34] (Table 2). The efficacies and potencies of the tested ligands were similar for Gq activation tested using the PKC biosensor versus direct activation with the Gα-Gγ biosensor (Table 1).

**Table 1.** Potencies (pEC50) and agonist-induced maximal response (“amplitude”) of G protein, protein kinase C and β-arrestin activation, normalised to the maximal response of Gs. Data are mean ± S.E.M of 2-7 independent experiments done either in triplicates or quadruplicates. Numbers in brackets indicate the fusion site of the luciferase in cases where different sensors were used. Please reduce the precision to the noise of measurement and move the tables into a separate file for the time being. They will need to be after references at submission. What is Emax – it always seems to be around 100. Do we need to report it?

| pathway | AVP |  | desmopressin |  | vasotocin |  | oxytocin |  |
| --- | --- | --- | --- | --- | --- | --- | --- | --- |
|  | pEC50 | response | pEC50 | response | pEC50 | response | pEC50 | response |
| **Gs (117)** | 8.76±0.08 | 100.9±2.2 | 8.39±0.14 | 92.8±4.3 | 8.08±0.11 | 85.8±3.4 | 6.89±0.16 | 86.8±3.6 |
| **Gi2** | 8.27±0.08 | 101.6±2.8 | 7.78±0.14 | 76.3±4.0 | 7.58±0.17 | 93.0±5.8 | 6.39±0.18 | 75.5±3.9 |
| **Gz** | 7.98±0.09 | 98.1±3.7 | 7.78±0.09 | 89.7±3.3 | 7.81±0.10 | 101.9±3.5 | 6.31±0.09 | 93.4±2.8 |
| **Gq** | 8.07±0.06 | 98.7±2.5 | 7.66±0.06 | 85.5±1.9 | 7.65±0.06 | 79.7±1.7 | 6.22±0.07 | 77.0±1.9 |
| **G12** | 7.79±0.21 | 86.2±7.2 | 7.81±0.25 | 111.4±9.2 | 7.69±0.22 | 125.2±10.7 | 6.10±0.16 | 118.5±6.4 |
| **G13** | 8.37±0.12 | 98.7±3.8 | 8.32±0.13 | 95.7±3.7 | 7.92±0.15 | 89.8±4.1 | 6.66±0.11 | 88.0±2.7 |
| **PKC (Gq)** | 8.69±0.07 | 99.7±2.3 | 8.01±0.10 | 88.8±3.4 | 8.06±0.16 | 98.9±5.5 | 6.25±0.11 | 84.4±3.6 |
| **PKC (G11)** | 8.79±0.09 | 99.1±2.9 | 8.03±0.09 | 94.4±3.2 | 8.28±0.12 | 92.1±3.8 | 6.48±0.09 | 97.5±3.0 |
| **PKC (G14)** | 8.63±0.19 | 97.3±6.6 | 7.50±0.43 | 79.4±18.6 | 7.38±0.24 | 142.6±14.0 | 5.83±0.25 | 96.2±9.6 |
| **PKC (G15)** | 9.26±0.08 | 99.5±2.2 | 8.43±0.1 | 109.2±3.2 | 8.49±0.09 | 113.2±3.3 | 6.77±0.14 | 98.1±4.4 |
| **β-arrestin 1** | 8.31±0.05 | 99.2±1.9 | 7.56±0.05 | 101.0±2.3 | 7.54±0.06 | 96.9±2.2 | 5.77±0.04 | 95.0±1.6 |
| **β-arrestin 2** | 8.23±0.05 | 99.1±2.0 | 7.50±0.04 | 98.8±1.7 | 7.45±0.05 | 93.9±1.8 | 5.84±0.04 | 95.1±1.7 |

**Table 2.** Dependence of log*B*_mm_ values on ligand amino-acid sequence, using AVP as the reference peptide. F3L: exchange of a phenylalanine at position 3 for a leucine, R8L: exchange of an arginine at position 8 for a leucine, R(L)8R(D) – exchange of the L-for D-arginine at position8.

| name | deami-<br>nation | F3<br>L | R8<br>L | R(L)<br>8<br>R(D) | $\log K_i$<br>[34] | Gs<br>$\log B_{\text{mm}}$ | Gi2<br>$\log B_{\text{mm}}$ | Gz<br>$\log B_{\text{mm}}$ | Gq<br>$\log B_{\text{mm}}$ | G12<br>$\log B_{\text{mm}}$ | G13<br>$\log B_{\text{mm}}$ |
| --- | --- | --- | --- | --- | --- | --- | --- | --- | --- | --- | --- |
| **AVP** | 0 | 0 | 0 | 0 | -8.97 | 0 | 0 | 0 | 0 | 0 | 0 |
| **desmopressin** | 1 | 0 | 0 | 1 | -7.51 | 0.01 | 0.08 | 0.05 | -0.1 | 0.1 | -0.1 |
| **vasotocin** | 0 | 1 | 0 | 0 | -7.6 | 0.02 | 0.01 | 0.02 | 0.11 | 0.1 | -0.02 |
| **oxytocin** | 0 | 1 | 1 | 0 | -5.81 | 0.17 | 0.17 | 0.05 | 0.13 | 0.1 | -0.05 |

**Figure 2.**
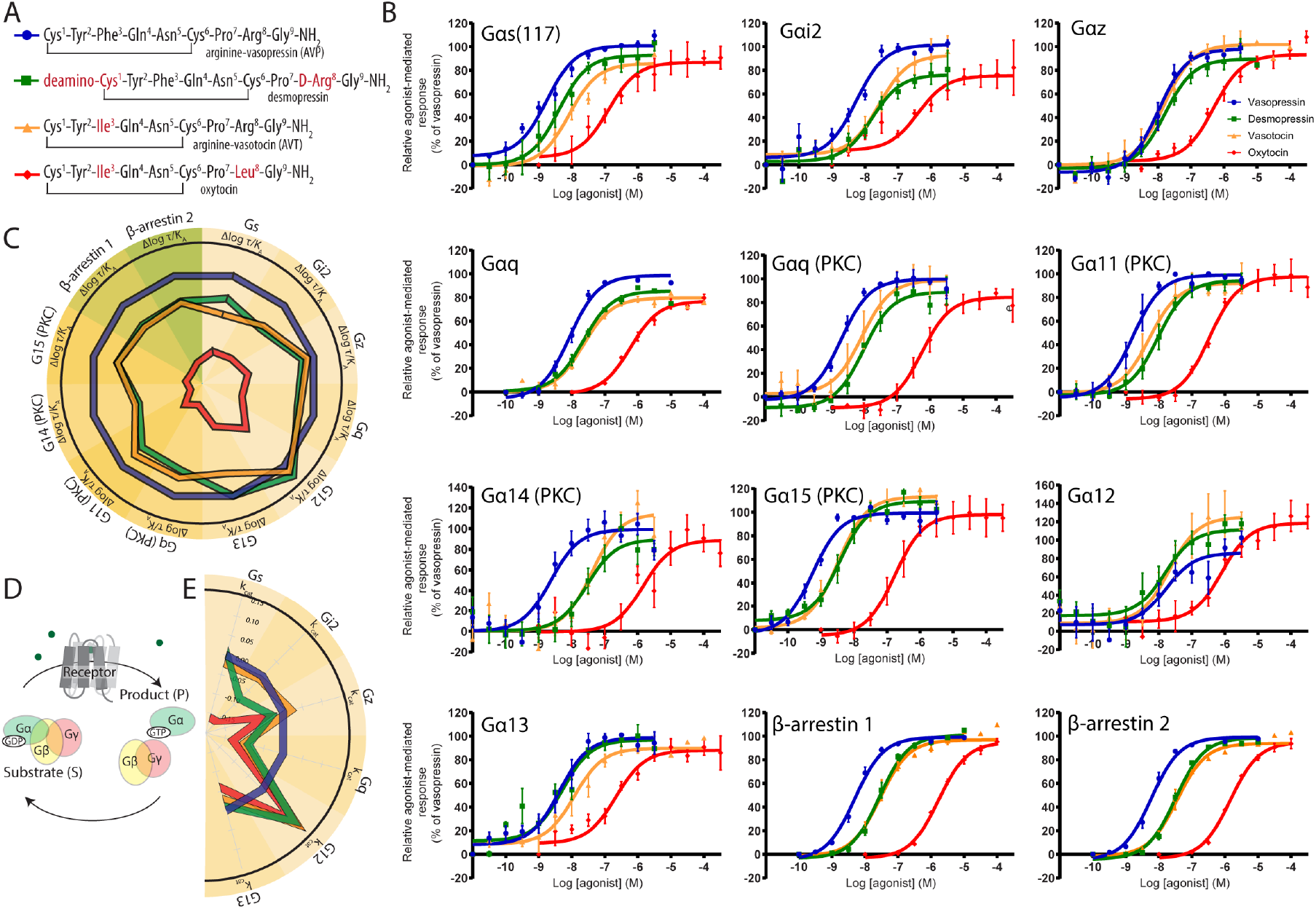
Biased signalling of V2R peptide ligands. **A.** The peptide ligands differ in only one or two amino acids (marked in red). **B.** Concentration-response curves of biosensor activation for all four peptides. **C.** Bias as calculated using the operational model for arginine vasopressin (AVP, blue), desmopressin (green), arginine vasotocin (yellow) and oxytocin (red). **D.** The schematic diagram of the Michaelis-Menten model of G protein activation. **E.** The Michaelis-Menten signalling bias.

We observed three different cases for the high-affinity ligands: First, equal potency for desmopressin and vasotocin, and a higher potency for AVP, which agrees well with the measured affinity values. This is the case for Gi2, Gq, G11, G14, G15, β-arrestin 1 and 2. Second, equal potency for all three agonists which we observed for Gz and G12. Finally, we observed that the ranked potencies for Gs and G13 were highest for AVP, medium for desmopressin and lowest for vasotocin for Gs; highest for AVP and desmopressin and medium for vasotocin for G13. In addition, we noticed that oxytocin shows a preference towards G protein engagement, while the potency for recruitment of β-arrestin is clearly lower. None of the ligands showed a preference for one of the two β-arrestins under the conditions of the study.

We calculated transduction coefficients (log(*τ*/*K*_A_) values) according to the operational model of agonism [24, 25, 35] to quantify ligand bias. AVP, the main agonist of vasopressin V2 receptor, was chosen as the reference agonist (Table 3). The choice of a reference agonist is necessary to eliminate observational and system bias [21]. The calculated transduction coefficients and their deviation from the reference agonist value (Δlog(*τ*/*K*_A_)) are shown in Fig. 2C. With the exception of Gz and G12, the ability of V2R to elicit response on all other signalling pathways was reduced by a factor of 3-5 for desmopressin and vasotocin. However, for oxytocin it was reduced by about a factor of 100, suggesting that it is a very poor agonist for V2R. On the other hand, comparison of the ability of the V2R to activate G proteins (Fig 2) relative to the reported affinity (Table 2) does not indicate that oxytocin is a particularly weak agonist in comparison to other peptides. Considering that all peptides were able to elicit the full amplitude of the biosensor response, the *τ* should be relatively large (i.e. >10) and the difference between the receptor-ligand complex formation and the effector activation response curve should be separated by at least a log unit [35]. On the other hand, the Δlog(*τ*/*K*_A_) values were significantly reduced for the lower-affinity ligands, implying a potential overestimation of the *K*_A_ value that reflects the affinity of the agonist for the “active” state of the receptor. This parameter is not the same as the one experimentally observed using ligand binding experiments which typically reflects the affinity of the effector-free receptor if an antagonist is used for determination of the *K*d. If an agonist is used to measure it, sometimes a bi-phasic binding curve and a high-affinity state is observed, reflecting an affinity of an agonist for the GPCR-effector complex [36–38]. The high-affinity stage is often observed under conditions of the effector protein present in abundance, however we observed no indications for the biphasic behaviour in our data (Fig 2), nor has it been reported for V2R in previous studies [34]. In the absence of the unusually high concentration of effector proteins, the receptor engages the G protein for a short time (e.g. **~** 100 ms) during the activation cycle, and is likely to spend most of its time “empty”. Shifting the receptor-ligand binding equilibrium takes minutes, given the concentration of the ligand and the kinetics of ligand binding. Therefore, the affinity of the ligand for the receptor is likely to be that of the uncoupled receptor. The apparent inconsistency of the activity analysis for low affinity ligands in comparison to high affinity ones, likely caused by the difficulties in estimating the “functional” affinity of the ligand for the receptor, prompted us to develop an alternative metrics that would link the bias calculations to the experimentally measured ligand affinity and would report changes in receptor activity towards a particular effector at an equal level of receptor saturation by the ligand.

**Table 3.**
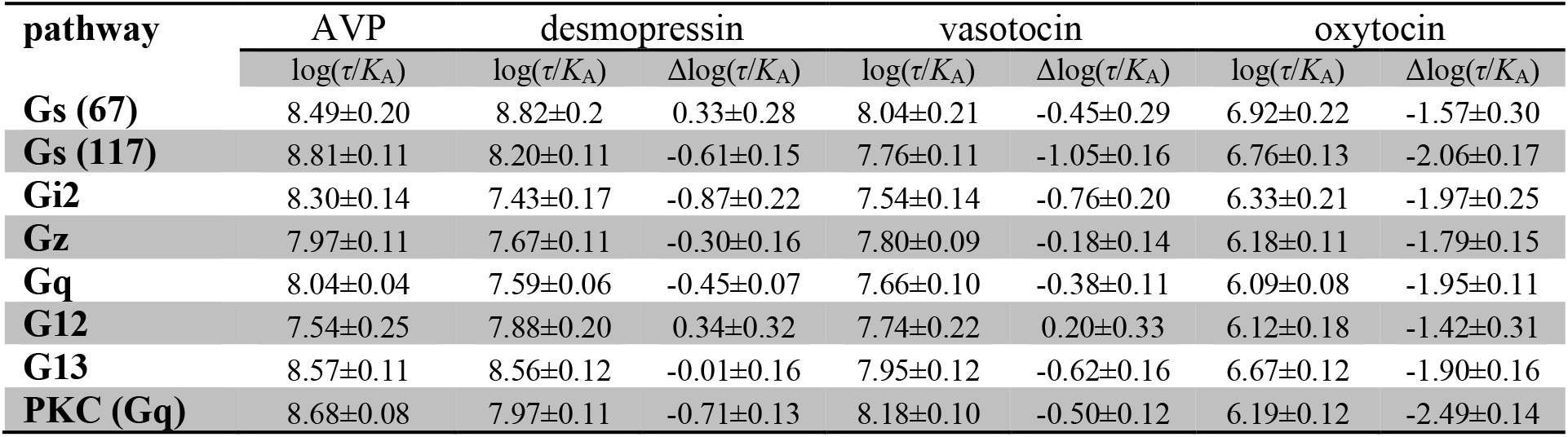
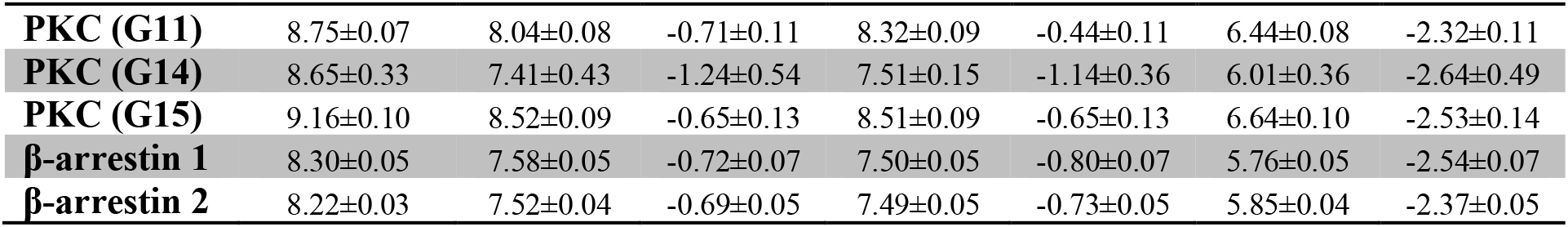
Transduction coefficients (log(τ/K_A_)) and Δlog(*τ*/*K*_A_) with AVP as reference ligand. Data are mean ± S.E.M of 2-7 independent experiments done either in triplicates or quadruplicates.

| pathway | AVP | desmopressin |  | vasotocin |  | oxytocin |  |
| --- | --- | --- | --- | --- | --- | --- | --- |
| | $\log(\tau/K_A)$ | $\log(\tau/K_A)$ | $\Delta\log(\tau/K_A)$ | $\log(\tau/K_A)$ | $\Delta\log(\tau/K_A)$ | $\log(\tau/K_A)$ | $\Delta\log(\tau/K_A)$ |
| **Gs (67)** | 8.49±0.20 | 8.82±0.2 | 0.33±0.28 | 8.04±0.21 | -0.45±0.29 | 6.92±0.22 | -1.57±0.30 |
| **Gs (117)** | 8.81±0.11 | 8.20±0.11 | -0.61±0.15 | 7.76±0.11 | -1.05±0.16 | 6.76±0.13 | -2.06±0.17 |
| **Gi2** | 8.30±0.14 | 7.43±0.17 | -0.87±0.22 | 7.54±0.14 | -0.76±0.20 | 6.33±0.21 | -1.97±0.25 |
| **Gz** | 7.97±0.11 | 7.67±0.11 | -0.30±0.16 | 7.80±0.09 | -0.18±0.14 | 6.18±0.11 | -1.79±0.15 |
| **Gq** | 8.04±0.04 | 7.59±0.06 | -0.45±0.07 | 7.66±0.10 | -0.38±0.11 | 6.09±0.08 | -1.95±0.11 |
| **G12** | 7.54±0.25 | 7.88±0.20 | 0.34±0.32 | 7.74±0.22 | 0.20±0.33 | 6.12±0.18 | -1.42±0.31 |
| **G13** | 8.57±0.11 | 8.56±0.12 | -0.01±0.16 | 7.95±0.12 | -0.62±0.16 | 6.67±0.12 | -1.90±0.16 |
| **PKC (Gq)** | 8.68±0.08 | 7.97±0.11 | -0.71±0.13 | 8.18±0.10 | -0.50±0.12 | 6.19±0.12 | -2.49±0.14 |
| **PKC (G11)** | 8.75±0.07 | 8.04±0.08 | -0.71±0.11 | 8.32±0.09 | -0.44±0.11 | 6.44±0.08 | -2.32±0.11 |
| **PKC (G14)** | 8.65±0.33 | 7.41±0.43 | -1.24±0.54 | 7.51±0.15 | -1.14±0.36 | 6.01±0.36 | -2.64±0.49 |
| **PKC (G15)** | 9.16±0.10 | 8.52±0.09 | -0.65±0.13 | 8.51±0.09 | -0.65±0.13 | 6.64±0.10 | -2.53±0.14 |
| **β-arrestin 1** | 8.30±0.05 | 7.58±0.05 | -0.72±0.07 | 7.50±0.05 | -0.80±0.07 | 5.76±0.05 | -2.54±0.07 |
| **β-arrestin 2** | 8.22±0.03 | 7.52±0.04 | -0.69±0.05 | 7.49±0.05 | -0.73±0.05 | 5.85±0.04 | -2.37±0.05 |

**Table 4.** Comparison of bias between pathways with Gs as a reference (ΔΔlog(τ/KA) values) and bias factors.

| pathway | desmopressin |  | vasotocin |  | oxytocin |  |
| --- | --- | --- | --- | --- | --- | --- |
| | $\Delta\Delta\log(\tau/K_A)$ | bias factor | $\Delta\Delta\log(\tau/K_A)$ | bias factor | $\Delta\Delta\log(\tau/K_A)$ | bias factor |
| **Gi2** | -0.26±0.27 | 0.55 | 0.28±0.26 | 1.93 | 0.08±0.31 | 1.21 |
| **Gz** | 0.32±0.22 | 2.07 | 0.87±0.21 | 7.46 | 0.27±0.23 | 1.86 |
| **Gq** | 0.17±0.17 | 1.47 | 0.67±0.19 | 4.69 | 0.11±0.21 | 1.28 |
| **G12** | 0.96±0.36 | 9.06 | 1.25±0.37 | 17.58 | 0.64±0.35 | 4.33 |
| **G13** | 0.60±0.22 | 3.99 | 0.43±0.23 | 2.69 | 0.16±0.24 | 1.44 |
| **PKC (Gq)** | -0.09±0.20 | 0.80 | 0.55±0.20 | 3.52 | -0.43±0.22 | 0.37 |
| **PKC (G11)** | -0.10±0.19 | 0.79 | 0.61±0.19 | 4.09 | -0.26±0.20 | 0.55 |
| **PKC (G14)** | -0.63±0.56 | 0.24 | -0.09±0.40 | 0.81 | -0.59±0.52 | 0.26 |
| **PKC (G15)** | -0.03±0.20 | 0.93 | 0.39±0.10 | 2.48 | -0.47±0.22 | 0.34 |
| **β-arrestin 1** | -0.11±0.17 | 0.78 | 0.25±0.17 | 1.76 | -0.48±0.18 | 0.33 |
| **β-arrestin 2** | -0.08±0.16 | 0.84 | 0.32±0.17 | 2.09 | -0.31±0.18 | 0.49 |

### Analysis of the G protein activation using the Michaelis-Menten formalism

One of the very promising approaches to describe and quantify the activity of GPCRs receptors *in vivo* and *in vitro* is by the enzymatic model [39, 40] and, in its simplified form, by the Michaelis-Menten formalism [41, 42]. In this scheme (Fig. 2D), the receptor is considered as an enzyme that catalyses the conversion of substrate to product, i.e. inactive G protein to activated G protein. In the simplest form (one-way activation of the G protein, upper part of the cycle presented in Fig. 2D), this will lead to eventual activation of all available G protein. Therefore, it is also important to consider that the activated G protein will be de-activated by alternative mechanisms, e.g. auto-hydrolysis of bound GTP to GDP. The balance between activation and deactivation will determine the concentration of the active G protein present that induces downstream signalling. The use of this model allows us to obtain the intrinsic enzymatic activity of the ligand-receptor complex towards a G protein that can be used for the calculations of intrinsic bias factors. To reliably fit concentration-response curves, it is essential to keep the model simple, with a minimal number of parameters. Therefore, the Michaelis-Menten formalism is preferred to the full enzymatic model as it has the same number of parameters as the operational model.

### In silico analysis of the Michaelis-Menten model of G protein activation

The first parameter is the concentration of available G protein (*S*_0_) which is related to the second parameter, the Michaelis constant *K*_m_. *K*_m_ describes the concentration of G protein at which the G protein activation by the receptor is half maximal. Another factor is the amount of active receptor, this is a combination of the receptor number and their activity (*R*_tot_·*k*_cat_), high activity can compensate for low receptor numbers and vice versa. The final parameter is the hydrolysis of GTP at the G protein which returns the G protein to its inactive state. The mathematical description of this model is included in the methods section (Eq. *1*-*3*). To explore and visualise the behaviour of the Michaelis-Menten model and the impact of variation of the parameters, we modelled effects of receptor activity, G protein deactivation, G protein concentration and the *K*_m_ of the G proteins towards receptor on the observed activation of the G proteins and, correspondingly, biosensor responses (Fig. 3 and 4). The system needs sufficient G protein (comparable to the *K*_m_ value or above) for the G protein activation to take place. However, further increase in the G protein concentration does not result in the increase of the potency of the response (i.e. left shift of the curve) as the response (under simulation conditions, see methods for details) follows the ligand binding curve (Fig. S2A). Correspondingly, *K*_m_ should be comparable or lower than the concentration of the G protein for the activation to happen (Fig. S2C). The system response is far more sensitive to the changes in the catalytic activity of the activated receptor (*k*_cat_) and the rate of the G protein deactivation (*k*_h_), compared to the changes in *K*_m_ or total G protein concentration *S*_0_. It must be noted that we combined the receptor density *R*_tot_ with the *k*_cat_ and modelled variations of *R*_tot_·*k*_cat_ parameter as this represents the activity of the receptor pool. The more active the receptor is, the fewer active receptor molecules are needed to reach 50% of the response, leading to a left-shift of the activation curve relative to the ligand binding. This is encouraging as we can capture the classical “receptor reserve” concept [43]. Contrarily, an increase in the rate of G protein deactivation *k*_h_ directly opposes the activity of the receptor (*k*_cat_). The faster the activated G protein hydrolyses bound GTP the harder it is to maintain an increased concentration of the active G protein.

**Figure 3.**
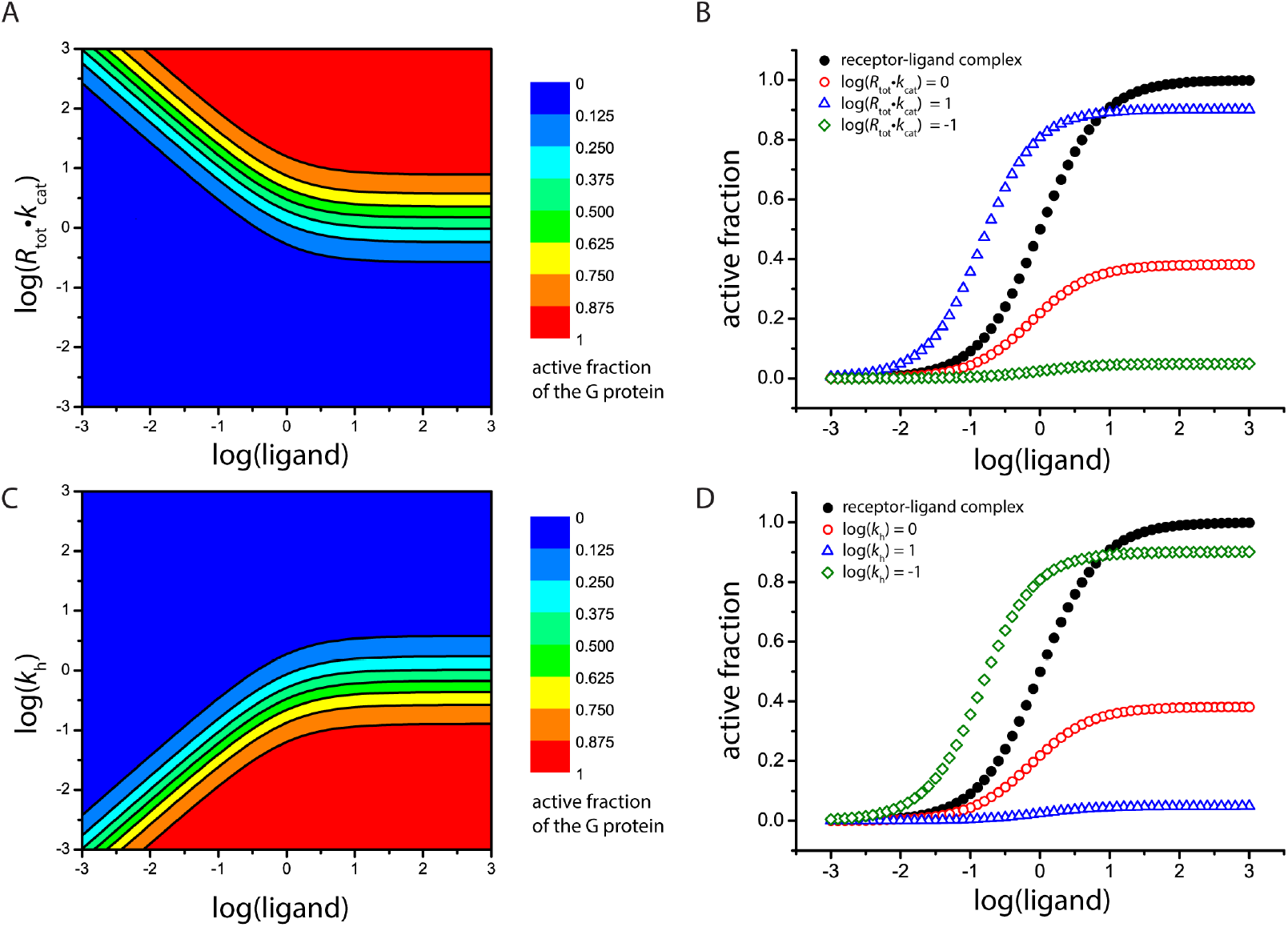
Activation of the G protein modelled according to the Michaelis-Menten formalism so that G protein activation is half maximal and log(ligand) = 0 refers to the ligand concentration where the concentration of ligand equals *K*_d_. *R_tot_* · *k_cat_* was modelled as one parameter because a high receptor number (*R_tot_*) can compensate for a slow *k_cat_* and vice versa. The units are arbitrary. (A) Fraction of active G protein as a function of receptor activity and amount (*R_tot_* · *k_cat_*). (B) examples of individual curves of (A) at several receptor activity levels. (C) Fraction of active G protein as a function of G protein deactivation rate constant *k*_h_. (D) examples of individual curves at different log(*k*_h_) values. Both parameters can result in a shift of the EC50 value as well as the amplitude of the response.

**Figure 4.**
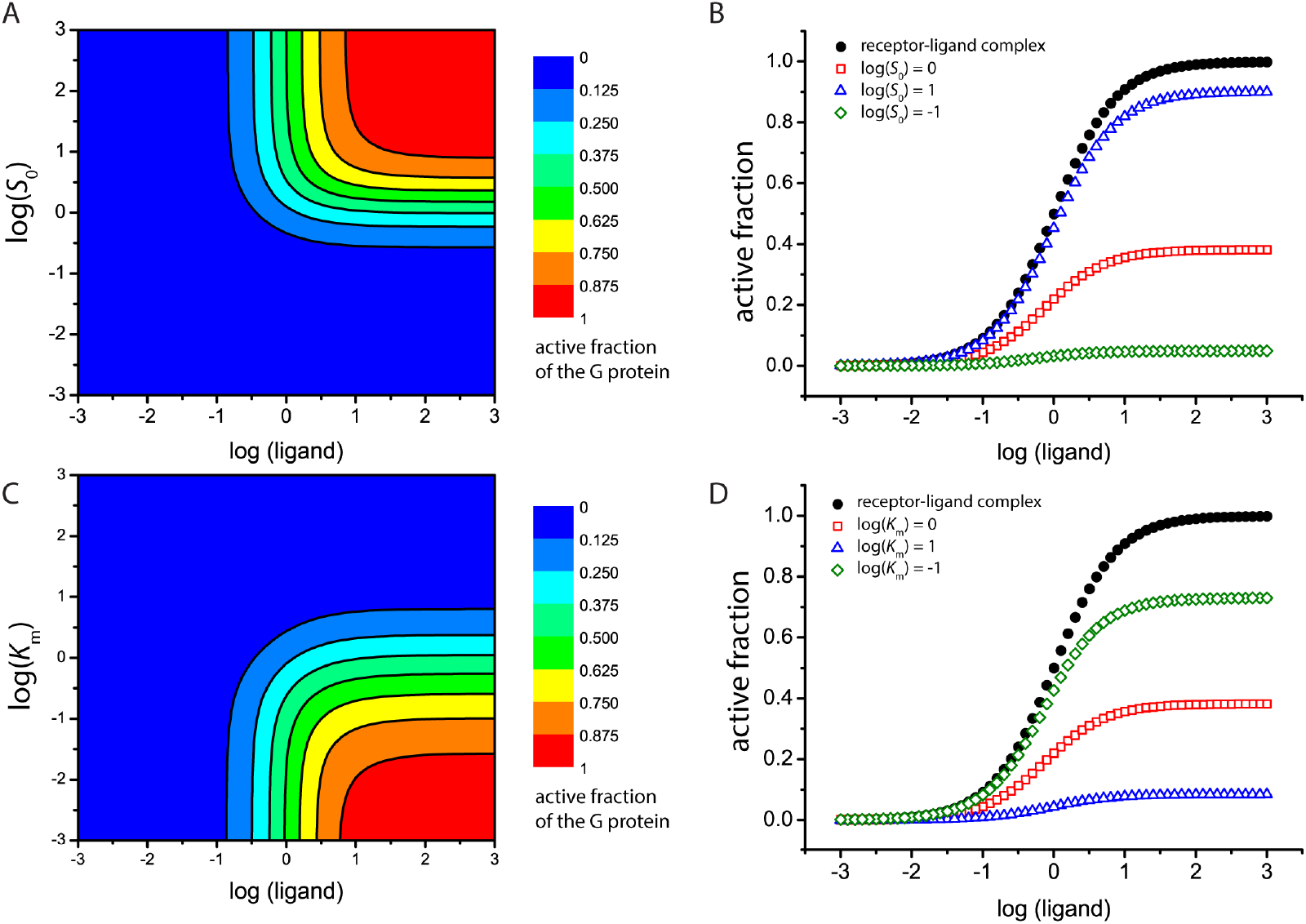
Dependence of G protein activation on total G protein concentration *S*_0_ and the Michaelis constant *K*_m_ for G protein–receptor interaction. Either *S*_0_ or *K*_m_ was varied, the other parameters were kept constant. In addition, the ligand concentration was varied over three orders of magnitude. (A) The fraction of active G protein is shown as a function of the total amount of G protein in the system (*S*_0_), assuming a *K*_m_ value of 1 (*S*_0_ = *K*_m_ at log(*S*_0_) = 0). (B) examples of individual curves at several G protein concentrations. (C) The fraction of active G protein as a function of the *K*_m_ value between the receptor and the G protein. The lower the value, the stronger the interaction. (D) Examples of the concentration-response curves normalised to the total amount of the G protein in the system. The EC50 value is not affected by these parameters, but the amplitude is.

### Application of Michaelis-Menten model to the experimental data

To apply this model to available experimental data, further assumptions need to be made. It is important to perform the experiments under the caveat of “all other things being equal”. The rate of the active G protein degradation is determined by the “system” (the cells we use for the experiments) and can be assumed to be constant. While we anticipate the *K*_m_ values for a given effector to be affected by the nature of the ligand, for a robust response it should still be comparable to the concentration of the G protein and has a limited potential to introduce significant error in the system. Therefore, the <u>*k*</u>_cat_ is the only significant parameter that determines the behaviour of the system, while all other parameters can be kept constant during the data analysis. As the values of these parameters are not known they are set to one in arbitrary units for the purpose of the fit and are cancelled out by normalisation. Normalisation of the of the obtained *k*_cat_ to that of a refence compound

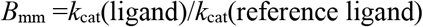

allows us to define a Michaelis-Menten bias factor *B*_mm_. A detailed explanation of the fitting procedure is presented in the methods.

We have applied this model to the BRET-based biosensor data presented above (Fig. 2B), as implemented in the DataFitter laboratory-developed software. Qualitatively, the results of bias calculations according to the Michaelis-Menten model (Fig. 2E) resemble those based on the operational model (Fig. 2C). The most noticeable difference is that the intrinsic activity and *B*_mm_ bias factors of oxytocin are comparable to those of other peptides (Fig 2D). This mirrors direct observations of the oxytocin activity as presented on Fig. 2B. The affinity of the ligand does not affect the calculated *k*_cat_ or *B*_mm_ (bias factors calculated according to the Michaelis-Menten model, see Methods) as opposed to the values derived from the operational model (Fig. 2C). As a result, the differences in the efficacy of the V2R towards G proteins in response to binding of diverse ligands are more accentuated. It is interesting to note that in comparison to AVP, all tested peptides have an increased ability to promote G12 engagement by V2R, but reduced ability to promote V2R-mediated activation of Gq. This parallels the analysis done using the operational model. In addition, oxytocin seems to have significantly reduced ability to promote V2R-mediated activation of Gs and Gi2 as compared to AVP.

Both operational model and Michaelis-Menten methods of calculating signalling bias have provided comparable results. However, Michaelis-Menten model provides more consistent estimates of the differences in signalling bias if the is a large difference in affinity of individual ligands. Taken together, these results strongly suggest that V2R signalling can be readily biased by ligands, and even relatively small structural differences in peptide ligand sequence seem to be sufficient to induce this effect.

## Discussion

### V2R promiscuity

Our data show that V2R can engage members of all G protein families, not only Gs and Gq as reported previously [10, 29]. A combination of our PKC activation data and previously published data showing an increase in cAMP levels after V2R activation, points towards activation of Gs and Gq by the V2R. All other G proteins (Gi2, Gz, G12, G13) are at least engaged. A recent paper showed that the V2R engaged G12 unproductively [31], which agrees with the engagement of G12 we observed. In addition, our analysis of the change of BRET signal in dependence of Gγ subunits (Fig. 1E) shows that G12 is the only G protein where we observed a decrease in BRET for Gγ1 but an increase in BRET for Gγ2 and Gγ5 suggesting that the interaction between the V2R and G12 is not a canonical activation, pointing to a different conformational change between Gα and Gβγ. We also confirmed a strong recruitment of both β-arrestins [32]. The biological significance of this promiscuity of engagement is beyond the scope of the present study but will warrant future studies. While V2R has a well-documented function in the kidneys where it controls water re-uptake, according to the Protein Atlas (www.proteinatlas.org) [44], it is expressed in practically all tissues, with the exception of the brain and the liver. Most Gα isoforms are also expressed in all tissues. Therefore, V2R has a possibility of interacting with all G proteins in native tissues, suggesting that its promiscuity may be biologically relevant. Dual Gs/Gi coupling has been reported for other receptors such as the β2- and the β1adrenergic receptors [45, 46] one possible rationale for simultaneous ctivation of Gs/olf and Gi/o proteins is to fine-tune the cAMP response. However, this does not account for the activation of Gq/11 and recruitment of G12/13 families. Another possibility is that the biological process triggered by the V2R may have to be mediated by a combination of signalling pathways, as each individual pathway may only induce a specific subset of cellular events. Recent medium- and large-scale profiling experiments confirmed that promiscuity is rather common among GPCRs [29, 47, 48].

### Ligand-induced signalling bias

The V2R is not only promiscuous in respect to the peptide ligands that activate it but its signalling is also biased by peptide ligands. All four peptides are very similar in sequence (Fig. 2A). They are constrained by a disulphide bridge and they likely adopt the same conformation and the same global binding pose in the V2R ligand binding pocket. Both the affinity of peptides for V2R as well as their signalling properties are affected by amino acid substitutions (Table 2). Any modification of the AVP, be it deamination at the N-terminus or substitutions at position 3 or 8 resulted in decreased ability to activate Gq and increased ability to recruit (most likely without activation) G12 (Fig. S1). In addition, double substitution of phenylalanine to isoleucine at position 3 and charged arginine at position 8 with hydrophobic leucine in oxytocin reduced the ability of V2R to engage Gs and Gi proteins relative to the other G protein subtypes engaged by the receptor. Similarly, substitution of the L-arginine for D-arginine at the same position in desmopressin relative to AVP may be responsible for the reduced Gi engagement compared to Gq. These specific effects of individual substitutions in AVP on peptide signalling properties imply that there must be hot spots in the ligand binding pocket for signalling bias that interact with the residues that were modified. However, in the absence of structural data on peptide binding it is difficult to say which amino acids in the ligand binding pocket are responsible for the signalling bias. Similar observations were reported for oxytocin receptor where small modifications of oxytocin peptide resulted in different signalling preferences [49]. Another extensively documented example are peptide ligands of angiotensin receptor [17, 50, 51]. Peptide ligand-induced biased has also been documented in chemokine receptors (reviewed in [52]). It is tempting to speculate that receptor-peptide pairs may have co-evolved as a mechanism to change the activity of ancestral receptors.

### Michaelis-Menten quantification of receptor activity

Previous work has shown applicability of the enzymatic model of GPCR activity [39, 40]. Kenakin and Christopoulos have commented on the apparent similarity between the operational model and the Michaelis-Menten equation [53]. However, despite the apparent mathematical similarity (both functions are hyperbolic), the actual solution for the steady-state concentration of activated G protein is rather different (see methods).

It should also be noted that the simplified Michaelis-Menten model presented here can describe G protein activation but not β-arrestin recruitment due to the underlying nature of the two processes. First of all, it assumes that the number of receptor molecules is small in comparison to the number of G protein molecules. For most receptors, even in the over-expressed systems, this condition is very likely to be satisfied. Secondly, it also assumes that the activation of the G proteins is non-reversible during the enzymatic step. Given the very high affinity of GTP to the G protein compared to GDP while their concentrations are comparable (0.1-0.5 mM) [54] and the slow hydrolysis-driven deactivation of G proteins, this condition is also very likely to be satisfied. While RGS proteins may control the rate GTP hydrolysis of Gi and Gq proteins, they only interact with active forms of Gα subunits after they have dissociated from Gβγ after the activation step [55]. Thirdly, it is important to consider the differences between the signals reported by the G protein, PKC and arrestin biosensors. While G proteins are activated directly by the receptors, PKC is activated by several nested enzymatic cycles, requiring a much more complicated model incorporating several Michaelis-Menten reactions. Thus, the data reflecting the activation of PKC do not lend themselves to the analysis using simple concentration-response curves as many more parameters are needed to define this system quantitatively. In contrast, arrestin biosensors report formation of the receptor-arrestin complex, with a 1:1 stoichiometry. As this is a binding rather than an enzymatic event, it would not be appropriate to analyse the results obtained with these arrestin-recruitment biosensors using Michaelis-Menten formalism. Recent reports suggested that arrestins may be activated and dissociate from the receptors while maintaining the active state, similarly to the G proteins [56]. If this is indeed the case, it may be possible to extend the use of this model onto arrestins. However, different biosensors directly reporting on the activation status of arrestin would have to be used [57–60]. One of the important advantages of the Michaelis-Menten formalism presented here is that it can be extended to describe the kinetics of signalling processes, and not only the steady-state equilibria. The appreciation that signalling may not be an equilibrium process and the importance of considering the kinetics in quantification of bias is growing [61]. We expect that as the biosensor and the data collection methodology improves and that kinetic data becomes routinely recorded, the application of Michaelis-Menten formalism would be of advantage in kinetic bias quantification.

## Conclusions

Our data show that V2R activates selected members of all G protein families and at least engage G12 in response to its native agonist AVP. However, closely related peptides that differ by only one or two amino acids or modification, show divergent signalling bias. This suggests that there are bias hotspots in the ligand binding pocket. We also present a method for quantifying the signalling bias based on Michaelis-Menten formalism that allows reliable separation of the intrinsic activity of the receptor/ligand complex from the effects of the affinity of the ligand to the receptor. Moreover, this method has a potential to be expended to quantify kinetic signalling bias in the future.

## Acknowledgements

This work was supported by the Swiss National Science Foundation grants 135754 and 159748 to DBV; Swiss National Science Foundation Doc.Mobility to FMH; and a Foundation grant (# 148431) from the Canadian Institute of Health Research (CIHR) to MB. BP was funded by a Fellowship Award from CIHR (2012-2015) and by a Fellowship Award from Diabetes Canada (2016-2018). MB holds a Canada Research chair in Signal Transduction and Molecular Pharmacology.

## Materials and Methods

### Vasopressin V2 receptor ligands

[Arg^8^]-Vasopressin (AVP) (Cys-Tyr-Phe-Gln-Asn-Cys-Pro-Arg-Gly-NH2; disulphide bridge: Cys^1^-Cys^6^, 1085.25 g/mol), Desmopressin acetate (deamino-Cys-Tyr-Phe-Gln-Asn-Cys-Pro-D-Arg-Gly-NH2; disulphide bridge: Cys^1^-Cys^6^, 1069.24 g/mol) and oxytocin acetate (Cys-Tyr-Phe-Gln-Asn-Cys-Pro-Arg-Gly-NH2; disulphide bridge: Cys^1^-Cys^6^, 1085.25 g/mol) were purchased from Genemed Synthesis Inc. (San Antonio, TX, USA) and [Arg8]-Vasotocin acetate (Cys-Tyr-Ile-Gln-Asn-Cys-Pro-Arg-Gly-NH2, disulphide bridge: Cys^1^-Cys^6^, 1050.22 g/mol) was from Sigma-Aldrich (Ontario, Canada).

### Biosensor constructs

Our biosensor measurements are based on BRETassay technology [62]. For the plasmids encoding for RlucII-Gα constructs, constructs were prepared using flexible NAAIRS linkers to insert Renilla luciferase (RlucII) into the coding sequence of human Gα versions. RlucII was inserted between amino acids Asp^94^ and Phe^95^ of Gαz using NAAIRSTRPRCT and TRPRCTNAAIRS as linkers. The Gαi1, 2 and 3 RlucII fusions contain a duplication of the respective loop where the RlucII was inserted; namely DSA and RLKIDFG for Gαi1, ADPS and NLQIDF for Gαi2, EAA and RLKIDFG for Gαi3, always followed and preceded by NAAIRS, respectively. Insertion positions were Gly^96^/Asp^97^ for Gαi1, Phe^95^/Ala^96^ for Gαi2 and Gly^96^/Glu^97^ for Gαi3. The GαoA construct was described previously (Richard-Lalonde et al., 2013). Gγ1, Gγ2 and Gγ5 were N-terminally tagged with GFP10 as described (Galés et al., 2006) and β-arrestin 1 and 2 were N-terminally fused to RlucII (Perroy et al., 2004).

### Cell culture and transfection

Human embryonic kidney (HEK) 293SL cells were transiently cotransfected with Flag-V2R, different RlucII-Gα variants, Gβ1 and GFP10-Gγ1 for G protein activation measurements and with Flag-V2R, RlucII-β-arrestin1 or 2 and CAAX-GFP10 for β-arrestin recruitment measurements. For Protein kinase C (PKC) activation, HEK293 ΔGq/11/12/13 cells were transiently co-transfected with Flag-V2R and unimolecular PKC biosensor for controls and with Flag-V2R, unimolecular PKC biosensor and either Gq, G11, G14 or G15 for activation experiments. The HEK293 ΔGq/11/12/13 cells were obtained by CRISPR-Cas9 technology [29]. Linear 25 kDa polyethyleneimine (PEI) (Polysciences Inc.) was prepared in phosphate-buffered saline (PBS) (Multicell) (PEI:DNA ratio 3:1). Per 0.24 million HEK293SL cells, 1 μg DNA was used. The cells were seeded into white Cellstar^®^ PS 96-well cell culture plates (Greiner Bio-One, Germany) at a density of 20,000 cells per well and grown for 48 h at 37°C with 5% CO2.

### Biosensor measurements

48 h after transfection the 96-well plates were washed with 200 μl PBS/well and 90 μl of Tyrode’s buffer (NaCl 137 mM, KCl 0.9 mM, MgCl_2_ 1 mM, NaHCO_3_ 11.9 mM, NaH_2_PO_4_ 3.6 mM, Hepes 25 mM, glucose 5.5 mM, CaCl_2_ 1 mM pH 7.4) were added and the cells were stored at 37°C with 5% CO_2_ for 2 h prior to the measurement. For the measurement the plates are incubated with 10 μl ligand or vehicle per well for 5 min, then 10 μl coelenterazine 400a (DeepBlueC) 2.5 μM final were added. After further 5 min of incubation, luminescence and GFP10 counts were measured at 410 and 515 nm, respectively, in a Synergy Neo (Biotek) plate reader using 0.4 s integration time.

### Preparation of ligands

All ligands were prepared in 0.1% (w/v) BSA, stock solutions were stored at −20°C while dilutions for the experiments were stored at 4°C. All ligand dilutions for experiments were used within 4 days of preparation.

### Data analysis

Data analysis was done in GraphPad Prism version 6.05 for Windows (GraphPad Software, La Jolla California USA, www.graphpad.com). All data points were normalised to the maximal response obtained with AVP and expressed as percentage. Values are given ±S.E.M for n experiments. Bias factors were calculated according to the operational model (Black and Leff, 1983; Gregory et al., 2010). The final equation used for non-linear curve fitting is:

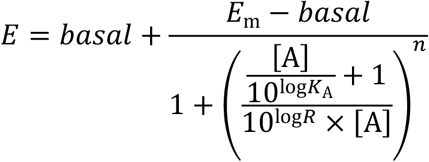

where *E* is the ligand effect, [A] is the agonist concentration, *E*_m_ is the maximal response of the system, basal is the signal in absence of ligand, *K*_A_ is the functional equilibrium constant, *R* is the transduction coefficient 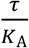where *τ* is an index for the efficacy of the agonist and *n* is the slope (Evans et al., 2011; Kenakin et al., 2012; Kenakin and Christopoulos, 2013; van der Westhuizen et al., 2014).

### Michaelis-Menten based description of the G protein activation by a GPCR

In the enzymatic model of GPCR activity, the G protein activation is catalysed by the receptor and is dependent on the agonist binding [39, 40]. The concentration of the active agonist-bound receptor *R*(*L*) is described by a binding isotherm:

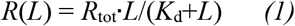

where *R*_tot_ is the total concentration of the receptor, *L* is ligand concentration and *K*_d_ is the ligand dissociation constant.

A minimal system (Fig. 2D) considers the formation of product (P, activated G protein, Gα-GTP) as a function of agonist-bound receptor concentration *R*(*L*), that is described by the Eq. *1* above, its catalytic activity rate constant *k*_cat_, as well as the Michaelis constant *K*_m_ for the G protein–receptor interaction. The rate of deactivation of the activated G protein (P) into inactive G protein (S) depends on the concentration of product and the hydrolysis rate constant *k*_h_ of GTP to GDP at the Gα subunit.

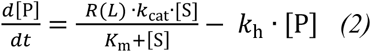

Considering the deactivation of the active G protein via GTP hydrolysis is an important feature of this model as it determines the concentration of the activated G protein.

At steady-state conditions there is an analytical solution yielding the concentration of activated G protein [P] as a function of the total (i.e., inactive and active combined) concentration of the G protein, *S*_0_.

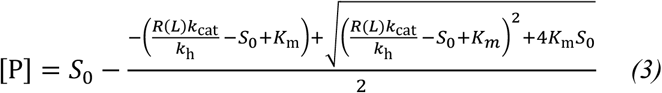

From the mathematical point of view, what matters for the steady-state solution is the apparent catalytic activity of the receptor–ligand complex in activating a G protein in a given system:

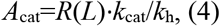

where *R*(*L*) and *k*_cat_ depend on the ligand affinity and concentration as well as the ligand signalling properties while *k*_h_ depends on the G protein and other system parameters.

Subsequently, the Eq. 3 could be simplified to

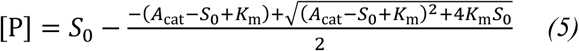

Correspondingly, the steady-state bias factor between two ligands for a given system can be expressed as

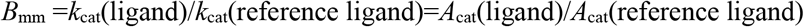

at the concentration of the ligand that results in the same occupancy of the receptor.

While in our experiments the values of *S*_0_ and *K*_m_ are not known, it is their value relative to the *A*_cat_ that would define the shape of the response curve. Therefore, for data fitting purposes we set their values to 1 and only interpret changes relative to the reference ligand. The data for individual ligands were fitted simultaneously to a MM model described above using the in house DataFitter software (D. Veprintsev).

### Simulations of Michaelis-Menten based description of the G protein activation by a GPCR

All simulations were performed using Cell Designer [63]. The value of the parameters of the system (*R*_tot_, *k*_cat_, *k*_h_, *K*_m_ and *S*_0_) were fixed to 1, while the value of the parameter presented on the Y axis and the ligand concentration were varied.

## Supplementary Figures

**Figure S1.**
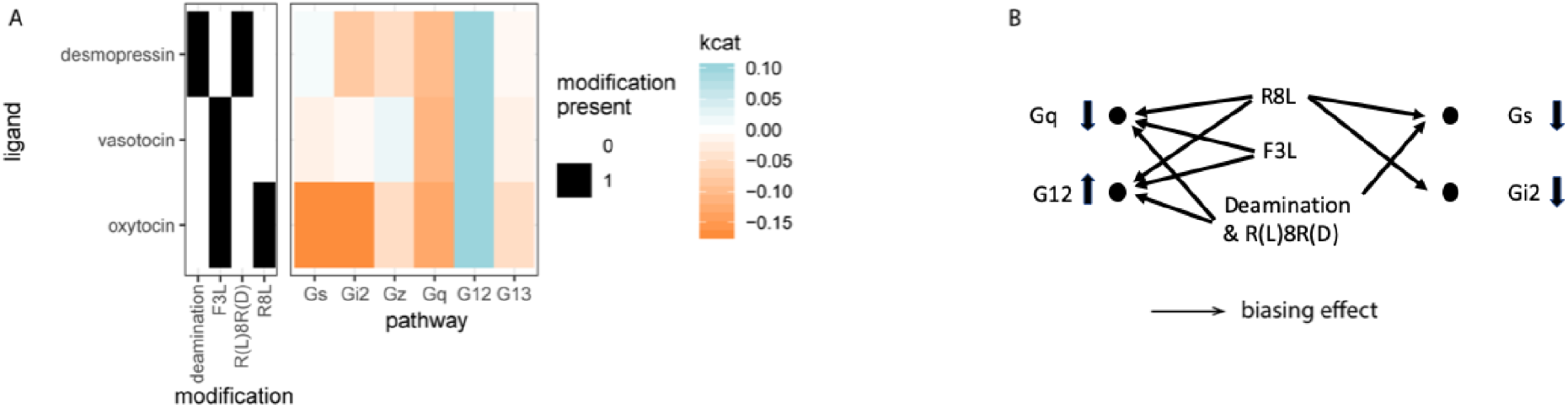
Correlation of the peptide modifications relative to the AVP with their signalling properties. **A.** A heat map showing the *k*_cat_ values. **B.** Observed trends of peptide modification on biased signalling.

